# Variance in cortical depth across the brain surface

**DOI:** 10.1101/2020.06.04.134593

**Authors:** Nick J. Davis

## Abstract

The distance between the surface of the scalp and the surface of the grey matter of the brain is a key factor in determining the effective dose of non-invasive brain stimulation for an individual person. The highly folded nature of the cortical surface means that the depth of a particular brain area is likely to vary between individuals. The question addressed here is: what is the variability of this measure of cortical depth? 94 anatomical MRI images were taken from the OASIS database. For each image, the minimum distance from each point in the grey matter to the scalp surface was determined. Transforming these estimates into standard space meant that the coefficient of variation could be determined across the sample. The results indicated that depth variability is high across the cortical surface, even when taking sulcal depth into account. This was true even for the primary visual and motor areas, which are often used in setting TMS dosage. The correlation of the depth of these areas and the depth of other brain areas was low. The results suggest that dose-setting of TMS based on visual or evoked potentials may offer poor reliability, and that individual brain images should be used when targeting non-primary brain areas.

## Introduction

Non-invasive technologies that read or that stimulate the brain, such as EEG, TMS, or tDCS, frequently make assumptions about the anatomical structure of the brain. This is necessary as, for most participants, the structure of the brain is not visible. Two common examples of this sort of assumption would be the use of a standard brain model in computing the source of dipole components in EEG analysis (e.g. Marin, Guerin, Baillet, Garnero, & Meunier, 1998), or the projection of a tDCS-derived computed electric field onto a standard brain model (e.g. Bikson, Rahman, & Datta, 2012). However to what extent can we rely on a standard brain image in understanding effects in an individual person? Are there particular regions of the brain where a standardised analysis is inadvisable?

The use of transcranial magnetic stimulation (TMS) is a good example of a technology that depends strongly on understanding the structure of the brain. In TMS, a brief electric current through a coil of wire induces a rapidly-varying magnetic field. This magnetic pulse can pass easily across the skull, and induces electric field changes in the brain tissues. This induced field is sufficient to cause action potential generation in neural tissue close to the focal point of the stimulation, and especially on the tops of the gyri (Bijsterbosch, Barker, Lee, & Woodruff, 2012). The so-called ‘hot-spot’ of the stimulation is thought to be around 1 cm^2^ in area at the surface of the cortex. The relatively small size of the hotspot, compared to the variance in the size and structure of the human head, means that a degree of accuracy is needed in targeting the stimulation.

Structural variance is one of many sources of variance in understanding the effect of TMS. Several studies have shown considerable intra- and inter-individual differences in responses to TMS. The motor evoked potential, or MEP, is an index of the excitability of the motor cortex, and is assessed by finding the motor hotspot, or the point on the scalp where a pulse of TMS induces a twitch in the participant’s contralateral hand. The functional target for this is thought to be the area of the primary motor cortex that projects to that hand (Rossini et al., 2015), although the precise anatomical location within Brodmann Area 4 is more variable (Ahdab, Ayache, Brugières, Farhat, & Lefaucheur, 2016). The motor threshold is defined as the stimulation intensity that causes a visible twitch in 50% of applied pulses (Rossini et al., 2015). Known sources of intra-individual variance in TMS-evoked potentials include the state of activation of the target brain area at the time of stimulation (Darling, Wolf, & Butler, 2006; Silvanto, Muggleton, & Walsh, 2008), muscular exercise (Liepert, Weiss, Meissner, Steinrücke, & Weiller, 2004) and wakefulness (Manganotti, Fuggetta, & Fiaschi, 2004). Sources of inter-individual variance include age (Matsunaga, Uozumi, Tsuji, & Murai, 1998; Rossini, Desiato, & Caramia, 1992), handedness (Triggs, Calvanio, Macdonell, Cros, & Chiappa, 1994) and many disorders of neural origin. There are not thought to be significant sex differences in MEP thresholds (Pitcher, Ogston, & Miles, 2003), although MEP latency is a function of distance between the cortex and the muscle, which differs between the sexes in most samples (Cantone et al., 2019). Intriguingly, MEP threshold seems to correlate between siblings, suggesting a degree of heritability in brain structure or excitability (Wassermann, 2002).

This paper will focus on the distance from the surface of the brain to the nearest point on the scalp surface. This is thought to be a crucial parameter for the efficacy of TMS, with modelling studies suggesting that the peak electric field relates to the depth from the surface of the cortex (Bijsterbosch et al., 2012). Experimental studies have determined that the required output of a magnetic stimulator increases by around 2.9% per millimetre of depth in order to have a meaningful effect (Stokes et al., 2005; Stokes et al., 2007). It is therefore important to know in dose-setting for TMS how deeply a target brain area lies from the scalp.

If all brains looked the same, it would be a simple matter to adjust stimulation intensity for a given area by scaling the stimulator output according to the depth of that area. However as well as the inter-individual differences in excitability outlined above, there are also a number of inter-individual differences in the morphology of the brain. The shape of the brain within the skull may be affected by many variables, including by dietary factors and hydration (Croll et al., 2018; Duning et al., 2005), by smoking (Gallinat et al., 2006), and by chronological age (Madan & Kensinger, 2018). This latter factor is of particular interest given the possibility of using non-invasive brain stimulation to address cognitive changes in healthy aging and in disorders of older age (Davis, 2017b; Perceval, Flöel, & Meinzer, 2016).

Without effective neuronavigation for target selection, most researchers might orient themselves on the participant’s head by finding either the motor or visual hotspot. The motor hotspot has been discussed already, while the visual hotspot is the point over the occipital cortex where stimulation induces an illusory visual sensation, or phosphene, that the participant can report. The optimum site for phosphene generation is around the inion, targeting the area of striate cortex that receives projections from around the centre of the visual field, where the phosphene is typically reported as a point flash of light (Cowey & Walsh, 2000; Kammer, 1999). These two hotspots differ principally in that motor effects are objectively measureable, while visual effects require a subjective report. But how to locate targets that lie away from these two primary areas, where the effect of stimulation is not readily obvious? Currently the best method is to acquire an image of the participant’s whole head, typically with an MRI scan, and to use this to manoeuvre a spatially-coregistered stimulator into position. This procedure, known as neuronavigation, means that individual brain and cranial anatomy can be taken into account, and allows the experimenter to target functionally-defined areas of the brain. Neuronavigation offers the most accurate means to target a given brain area such as the motor hotspot (Sparing, Buelte, Meister, Paus, & Fink, 2008), although even this method may not always precisely locate the optimum location for inducing motor effects (Herwig et al., 2002). Neuronavigation offers the best available method for precise delivery of TMS to a target area, although it is still prone to imprecision and artefacts (Julkunen et al., 2009; Säisänen et al., 2015).

Alternative methods of locating a stimulator include measuring the head and marking points that lie on intersections of the lines of latitude and longitude. This latter method is the principle behind the International 10-10 and 10-20 systems, commonly used in EEG studies (Jasper, 1958). In these systems, four key points are identified on the head (nasion, inion, and left and right pre-auricular points) and the ‘equator’ is defined as the line that runs through all these points. Following this, lines can be connected between pairs of points, such as the midline, which connects the nasion and inion, and further points can be derived such as Cz, which is where the point where the midline crosses the line which connects the two pre-auricular points. Cz is often used as a (so-called) ‘control site’ for TMS studies as it sits at the top of the head, and it is often suggested that there is no brain tissue under this point (Davis, Gold, Pascual-Leone, & Bracewell, 2013). Other derived points include F3 and F4, which lie bilaterally above the prefrontal cortex and are often used in procedures that aim to affect mood states (Austin et al., 2016). A second alternative method might be simple measurement of distances. For example the premotor or dorsolateral prefrontal areas may be located by measuring a fixed distance anterior to the motor hotspot; however this method is considerably less accurate than other systems (Ahdab, Ayache, Brugières, Goujon, & Lefaucheur, 2010).

With the increase in availability of brain stimulation techniques, many researchers have pointed out the difficulty of determining a dose of stimulation. In other therapeutic contexts, such as paediatric pain medication, there may be a simple metric that determines the correct dose of analgesic, such as a figure expressed in milligrams per kilogram of bodyweight. No such metric exists for NIBS. Indeed the number of available factors means that a wide array of parameters must be reported in order to make an intervention reproducible (Peterchev et al., 2012). These parameters include the location of the stimulation, the intensity, the duration, and the dynamic pattern of the stimulation. However a number of person-specific factors also apply, as mentioned already.

One key factor that affects the physiological response to NIBS is the physical structure of the cortex underlying the coil or pad. TMS appears to generate action potentials at the ends of axons, particularly in pyramidal cells in layers 3 and 5 of the cortex (Aberra, Wang, Grill, & Peterchev, 2020; Seo, Schaworonkow, Jun, & Triesch, 2016), and is dependent on the orientation of the changing electric field relative to the tissue (Salvador, Silva, Basser, & Miranda, 2011). This hints at the sensitivity of the effects of TMS to the precise structure of the brain: if the coil is not aligned correctly to the target tissue, a higher stimulator output will be required to generate an effect. Brain morphology varies between participants in many measures, including gyrification, spatial complexity and gross size (Madan & Kensinger, 2018). It is therefore important to understand how morphological variance might lead to variance in the effects of NIBS.

### Aims and hypotheses

The first aim of this study is to determine the relative variance across the cortex of the scalp-to-cortex distance. We would expect individual points in the brain to vary in depth, due to the folded nature of the cortical sheet. However the specific question asked here is whether these depths differ in their coefficient of variance (CoV; standard deviation divided by mean). The null hypothesis would be that the CoV is constant across the cortical surface; any deviation from this would be evidence that a given region has unusually high or low variance between participants. The variance will be estimated for each point in a normalised space for the whole surface, and will be analysed separately for five regions of interest.

The second aim is to determine which areas of the cortex have a strong correlation in depth with the hand area of primary motor cortex and with the foveal area of primary visual cortex, which are used in setting TMS dosage for non-primary brain areas. These primary thresholds may correlate with each other under carefully-controlled titration procedures (Deblieck, Thompson, Iacoboni, & Wu, 2008), although several studies find poor correlation, possibly due to a relatively high variance in the phosphene measure (e.g. Stewart, Walsh, & Rothwell, 2001). The use of a large set of brain images will allow for the investigation of intra-individual variance.

## Materials and methods

### Brain images

MRI images were taken from the Open-Access Series of Imaging Studies (OASIS) database (https://www.oasis-brains.org/; Marcus et al., 2007). Of the 436 scanning sessions in the full OASIS data set, only scans from participants aged 25-60 (inclusive) were selected. Where a participant was scanned twice for quality analysis purposes, only the first of the two sessions was included here. This left 111 unique sessions, contributed by 65 female and 46 male participants, with a mean age of 41.7 (SD 11.7). During analysis, images of 17 of these participants were removed, meaning that the final analysis was conducted on 39 male and 55 female participants, with a mean age of 40.95 years (SD 11.47). All participants were right-handed, and where a Clinical Dementia Rating (CDR) was available, all scores were zero (meaning no evidence of dementia: Morris, 1993).

Image acquisition is described in full by Marcus et al. (2007). Briefly, the images were acquired on a 1.5 Tesla Siemens Vision MR scanner, with an MP-RAGE sequence optimised for good contrast in anatomical imaging (TR=9.7ms, TE=4.0ms, Flip angle 10°, 1×1×1.25mm resolution). Each participant was scanned four times within a session. The OASIS data set contains a number of raw and processed images, including the four anatomical scans, plus an average image of these four scans. For the present analysis only the subject-specific averaged anatomical image was selected, with all further processing conducted using custom Matlab scripts based on scripts available in SPM12b. Scripts written for the present analysis are available at https://github.com/nickjdavis/ScalpGM.

### Preprocessing

The anatomical scans were first rotated to approximately the same alignment as MNI-format brain images, using a custom script (oasis_reorient.m written by Ged Ridgway). Following this, the SPM toolbox was used to segment the images into different tissues types, of which only the grey matter and the soft tissue classes will be discussed further.

### Distance analysis

The soft tissue class includes the tissue of the scalp. A custom Matlab script, using the built-in convhull function, was used to fit a 3d convex hull to the outside of this tissue class. This was assumed to represent the scalp surface, or the closest that a non-invasive coil or electrode could be placed to the brain. With this convex hull, it was then possible to compute, for each point in the grey matter image, the Euclidean distance in millimetres between that point in the brain image and the points of the convex hull. The minimum distance between each point in the grey matter and the scalp was saved into a new image, called the distance image. In order to compare distance profiles across participants, the images were warped (using the SPM function) into the MNI template. This meant that for each participant, we obtain an image in standard (MNI) space that represents how far (in native space) that point originally lay from the scalp. Each distance image was visually inspected at this point, and images that were obviously misaligned were removed from the analysis. 17 images were removed for this reason, leaving 94 images for further analysis.

For statistical analysis, three aggregate images were created. The first was the mean distance at each point in MNI space for all images. The second was the standard deviation (SD) of each of these points. Finally, the coefficient of variation (CoV) was calculated as a pointwise division of the SD by the mean. Since SD scales with the mean for normally-distributed data, the CoV more clearly shows areas of high variance in depth. The AAL template (Tzourio-Mazoyer et al., 2002) was used to identify the regions of the MNI-space depth image corresponding to five commonly-stimulated brain regions: the precentral gyrus (containing the primary motor cortex); the superior occipital gyrus (containing the primary visual cortex); the middle frontal gyrus which is involved in language and attention processes (e.g. Japee, Holiday, Satyshur, Mukai, & Ungerleider, 2015; Wen et al., 2017); the angular gyrus of the parietal cortex; and the superior temporal gyrus (containing the primary auditory cortex). For each of these areas, the mean and standard deviation of depth was calculated for the left and right hemispheres separately. Mean depth was subjected to a repeated-measures analysis of variance with factors for Area and for Hemisphere. The mean and standard deviation were used to calculate CoV, which was defined as the voxelwise SD divided by the voxelwise mean depth.

Although CoV is primarily a descriptive measure, it obeys a known statistical distribution, and is therefore amenable to hypothesis tests (Argaç, 2005). While there is no commonly-agreed ‘good’ level of CoV (as compared to, say, levels of effect sizes: Cohen, 1992), we can at least test CoV values against a target value, using a version of Student’s t-test (Doornbos & Dijkstra, 1983). For the present work a CoV value of 0.2 has been somewhat arbitrarily taken as a reference point; this value accords with other values in the biological literature (e.g. Pélabon, Hilde, Einum, & Gamelon, in press). It is worth noting that when CoV is 0.2, 95% of observations lie within 39.2% of the mean, which would seem to be rather a large margin of error for many applications (Van Belle & Martin, 1993).

A second set of images was created that mapped the correlation between the depth of the two key areas and the rest of the brain surface. These key areas were the primary motor cortex (M1) and the primary visual cortex (V1). To create these maps, a region of interest was defined as a sphere of 8 mm radius, centred on MNI positions within these areas. The motor ROI was centred on MNI coordinates [-37, -25, 62], namely the left primary motor cortex (Mayka, Corcos, Leurgans, & Vaillancourt, 2006). The occipital ROI was determined as the pole of the occipital lobe of the left hemisphere, at MNI coordinates [-12, -100, 5]. The depth of these regions were then each correlated with the depth of each voxel in the depth image for each participant. Images for the correlation coefficient (Pearson’s *r*), for the number of voxels included, and for the uncorrected p-value were saved for analysis.

Statistical analyses were conducted in SPSS version 25, with an alpha level of 0.05. Terms in the analyses of variance that violated Mauchly’s test of sphericity were corrected using the Greenhouse-Geisser method.

Figure 1 shows an example of the processing pipeline, in 2d cross-section.

**FIGURE 1.**
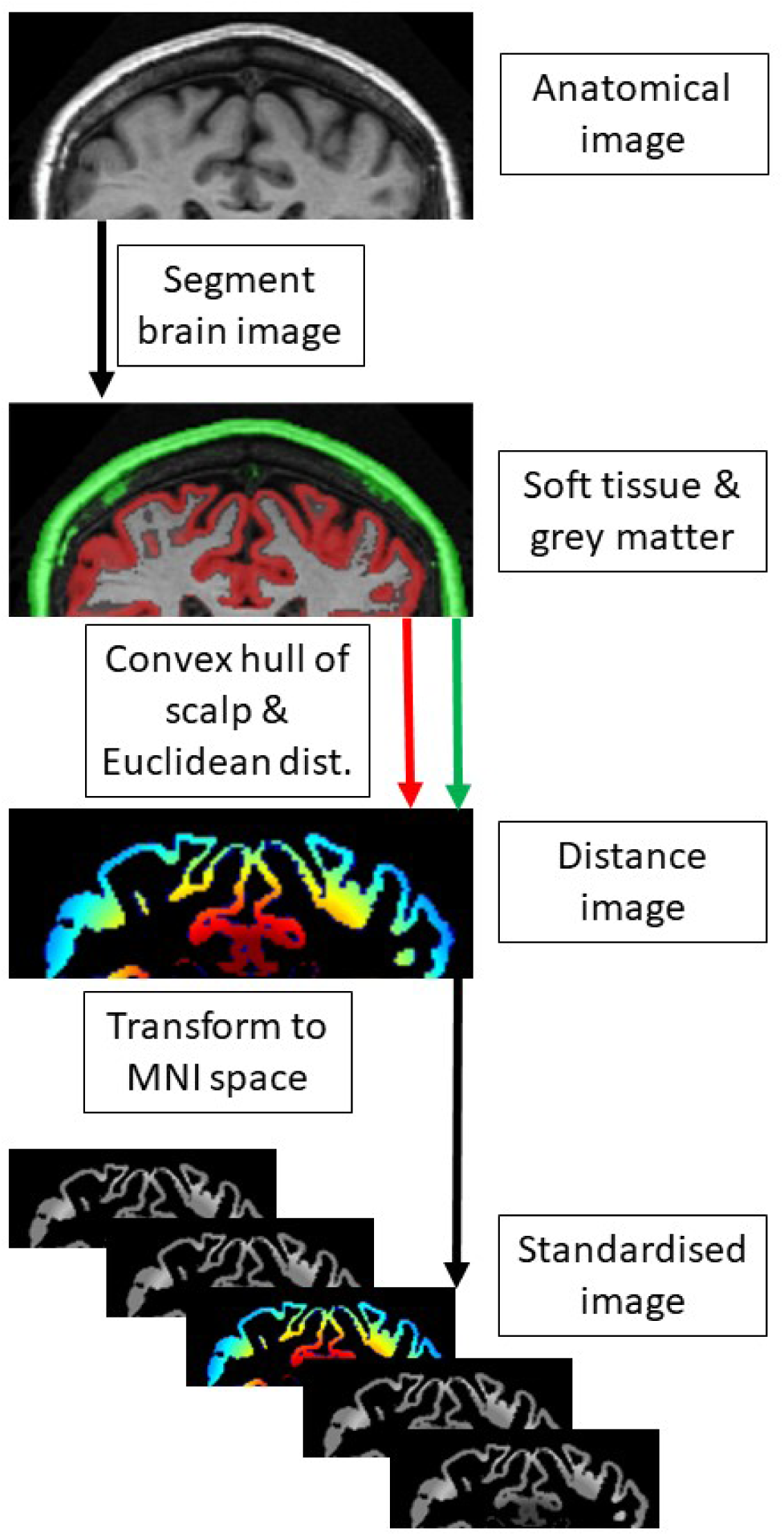
Processing stages. The analysis begins with an anatomical image of the brain, which is segmented to recover the soft tissue and grey matter. A convex hull is fitted to the soft tissue, and from this a map of minimum scalp-to-grey matter distance is obtained. This image can then be warped to standard (MNI) space, and combined with other images for statistical analysis.

## Results

### Effects of chronological age

The OASIS data sheet contains estimates of total intracranial volume (eTIV) and the proportion of intracranial voxels that contain brain tissue (normalised whole-brain volume, nWBV). In the sample used here, eTIV did not correlate with age [r(92) = -0.12, p=0.271], however nWBV did correlate significantly with age [r(92) = -0.51, p<0.01]. Linear regression suggested that brain volume decreased by approximately 0.001 litres per year [F(1,92) = 31.55, p<0.001; R^2^ = 0.247]. These results agree at a qualitative level with the results for the full sample as presented by Marcus et al. (2007), and are illustrated in Figure 2.

**FIGURE 2.**
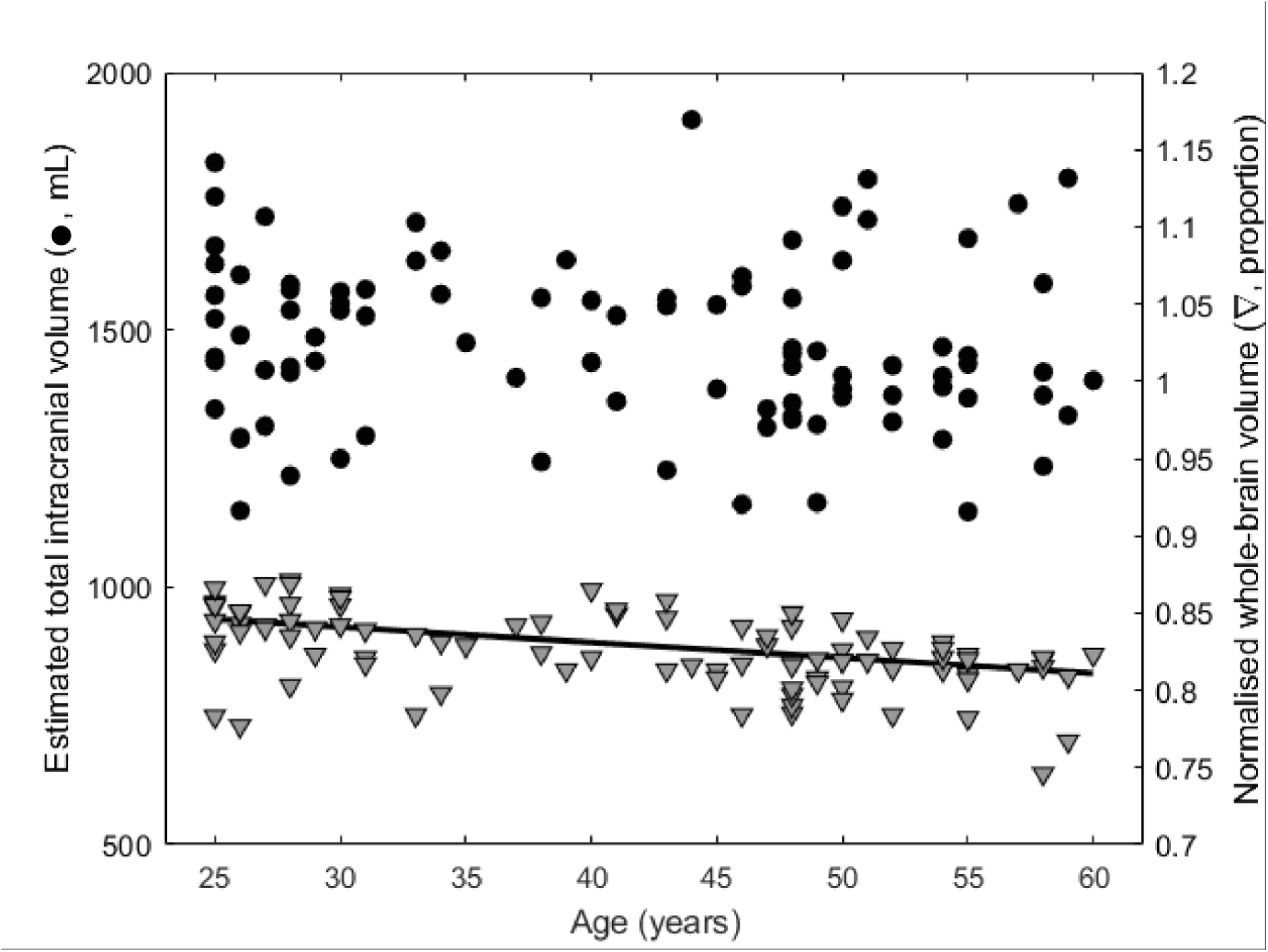
Scatterplots of total intracranial volume (eTIV) and normalised whole-brain volume (nWBV) against age. eTIV does not change through the adult years, however nWBV declines steadily at a rate of around 1 mL per year.

The depth of the M1 ROI correlated significantly with age [r(92)=0.2471, p<0.05], however this was not true for the V1 ROI [r(92)=0.0893, p=0.392]. The depth of the M1 ROI did not differ between male and female participants [t(92)=0.891, p=0.376], but the depth of V1 did differ [t(92)=2.860, p<0.01].

### Mean depth of regions of interest

Figure 3A shows the mean depth of the cortical surface, projected onto an inflated map. The standard deviation of this depth (Figure 3B) reveals areas of high and low variance. However the standard deviation of a variate scales with the mean. So the coefficient of variation (CoV), illustrated in Figure 3C, provides a scale-free measure of the relative variance of the depth of the cortical surface. Figure 4 shows the regions of Figure 3C where the CoV is significantly higher or lower than an arbitrary value of 0.2 (calculated using a modified and uncorrected pointwise t-test). Although few areas are lower in CoV than 0.2, several regions are significantly greater than this value, including the dorsal ends of the central and postcentral sulci, and lateral areas of occipital and temporal lobes.

**FIGURE 3.**
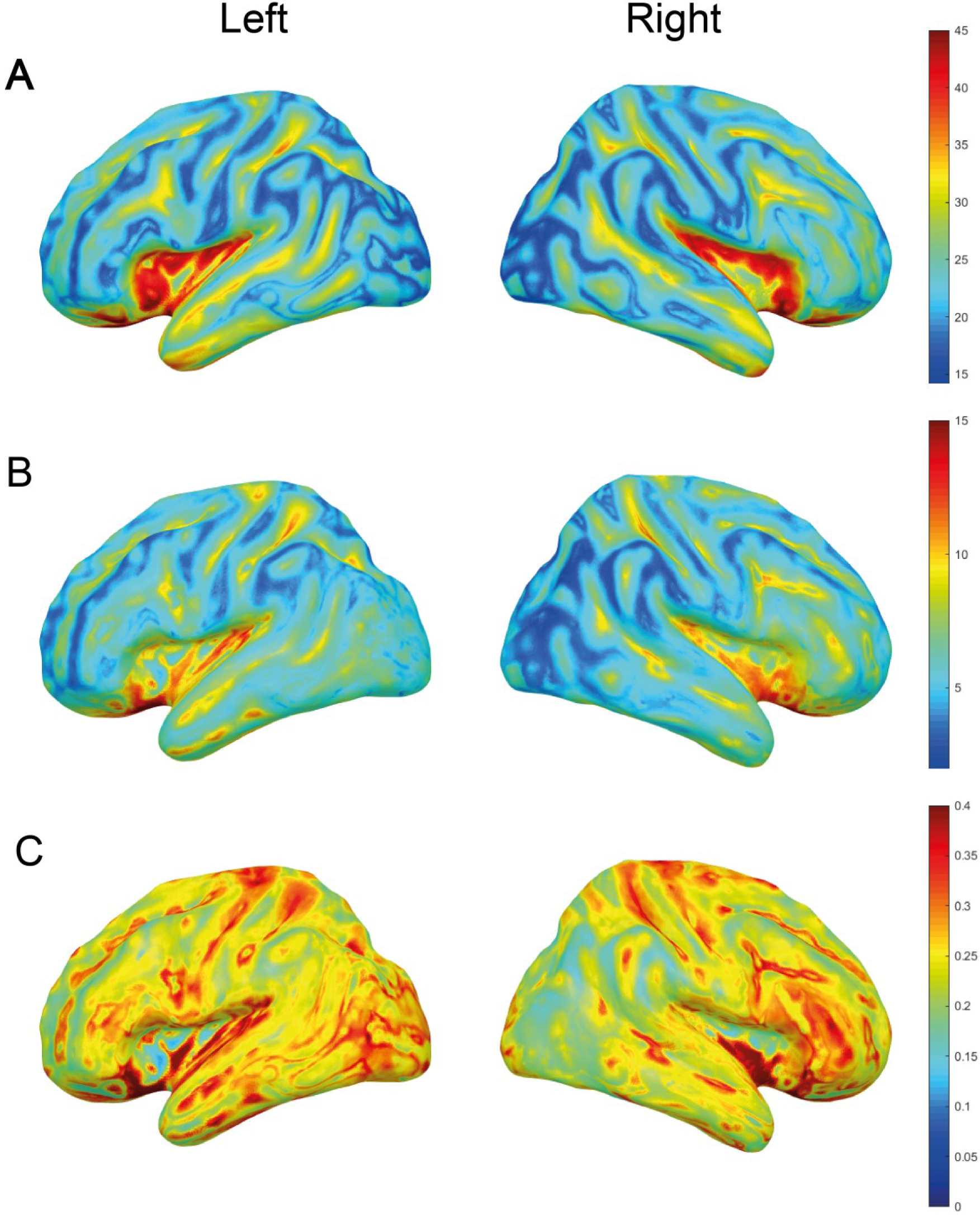
Group average images for the mean depth (A) in millimetres, the standard deviation of the mean depth (B, millimetres) and the coefficient of variation (C). The lateral surfaces of the left and right hemispheres are shown in an inflated projection.

**FIGURE 4.**
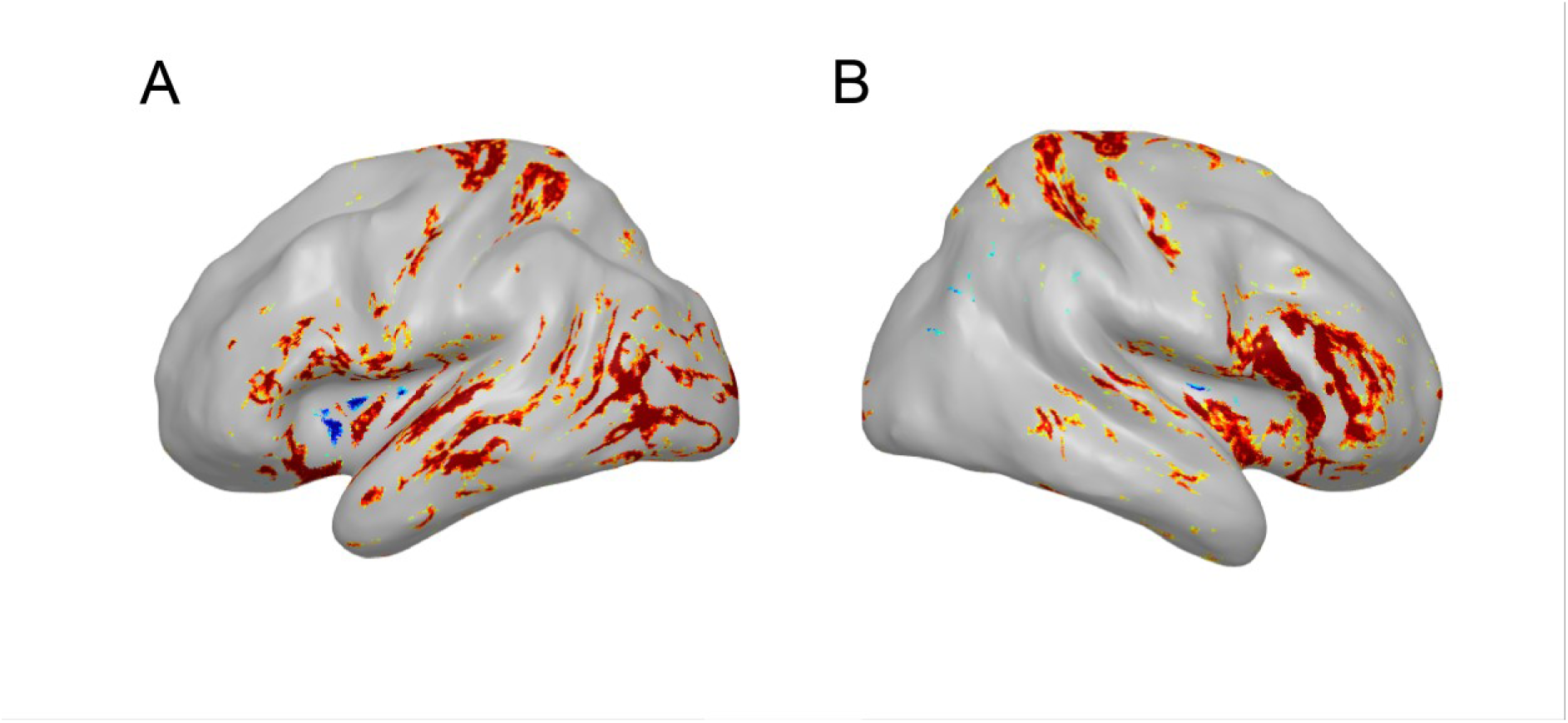
Regions of the coefficient of variance map shown in Figure 3C which differ significantly from 0.2. In this image few regions have a lower CoV than 0.2, but many functionally-important regions of the lateral cortex are significantly higher.

The AAL atlas was used to find the voxels in each normalised brain image that corresponded to defined brain regions. These regions were the precentral gyrus, the superior occipital gyrus, the superior temporal gyrus, the middle frontal gyrus and the angular gyrus. Figure 5 illustrates the mean and standard deviation of the subjectwise mean depth of these areas for the left and right hemispheres, as well as the coefficient of variation of these depths.

**FIGURE 5.**
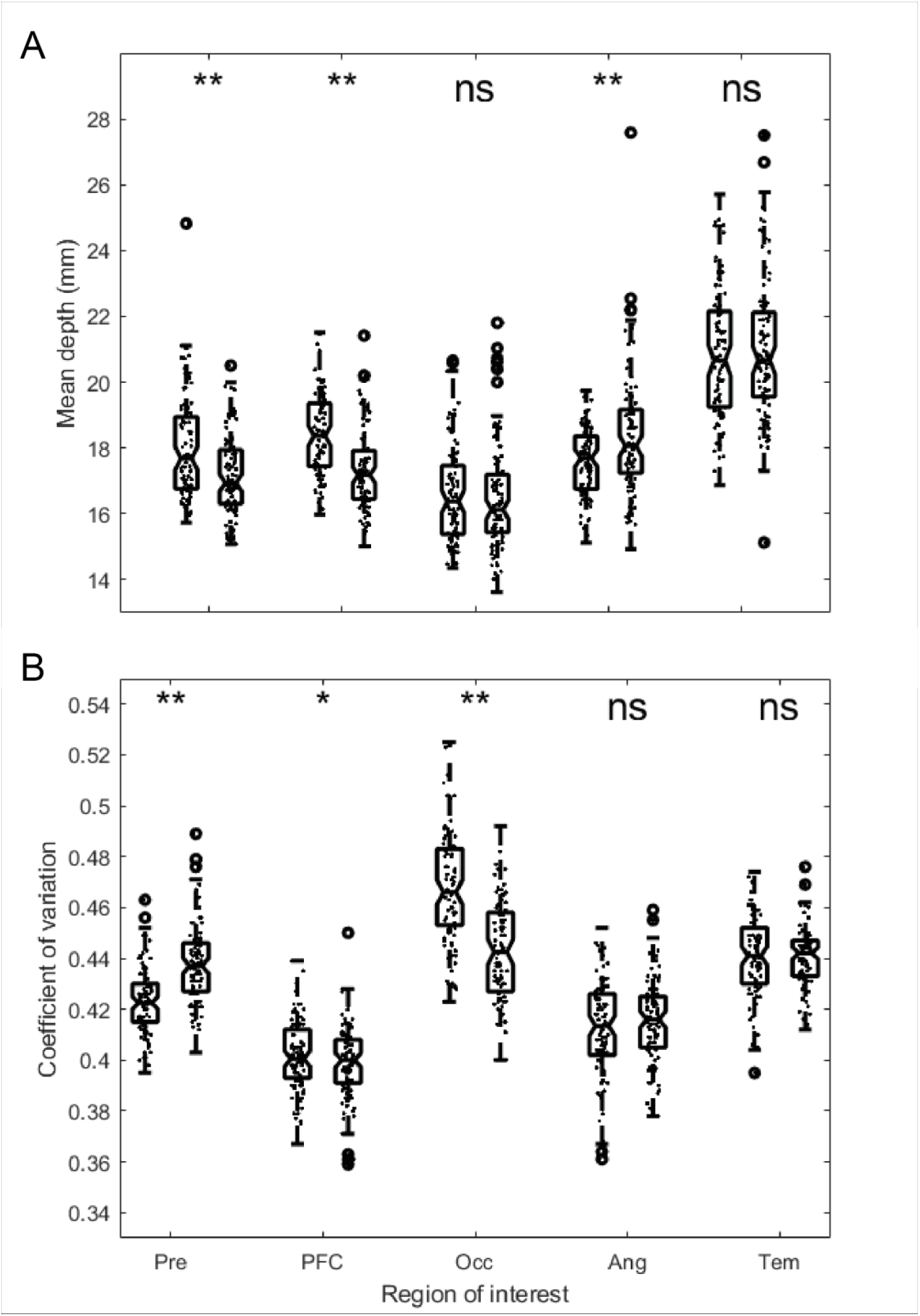
Mean and coefficient of variation (CoV) in each of five regions of interest (ROIs), for left and right hemispheres separately. Mean depth (Figure 5A) varies across the ROIs, as might be expected, and this depth differs between the hemispheres in the precentral gyrus (Pre), the middle frontal gyrus of the prefrontal cortex (PFC) and the angular gyrus (Ang), but does not differ between the hemispheres for the superior occipital gyrus (Occ) or the superior temporal gyrus (Tem). For the CoV, in Figure 5B, we see a different pattern, with the precentral gyrus, the middle frontal gyrus of the prefrontal cortex and the superior occipital gyrus showing different relative variance between left and right, but this does not differ for the angular gyrus (Ang) or the superior temporal gyrus (Tem). * p<0.05, ** p<0.01.

As expected, mean depth varied across these five brain areas [F(2.93,272.36)=224.35, p<0.001, eta^2^=0.707], with the temporal cortex the deepest of these areas, and the occipital region the shallowest. Overall depths were not significantly different between left and right hemispheres [F(1,93)=8.80, p=0.051]. However these effects interacted [F(2.71,252.39)=23.91, p<0.001, eta^2^=0.205], meaning that the relative depth of each brain area differed between left and right hemispheres. Depth data for these ROIs is shown in Figure 5A. The equivalent analysis performed on CoV scores revealed a superficially similar pattern of statistical results, with a significant main effect of area [F(2.98,277.38)=331.80, p<0.001, eta^2^=0.781] and a marginally significant effect of hemisphere [F(1,93)=5.66, p<0.05, eta^2^=0.057], and again an interaction between the terms [F(2.89,268.83)=66.29, p<0.001, eta^2^=0.416]. However, as shown in Figure 5B, the distribution of results is somewhat different, with the prefrontal region showing the lowest amount of CoV, and the occipital region showing the greatest.

### Correlations between regions

Figure 6 illustrates the correlation coefficient of each point of the cortex with the depth of the motor cortex (Figure 6 A, B) and with the visual cortex (6 C, D). Correlation coefficients are low across the cortex, and nowhere do they exceed 0.65.

**FIGURE 6.**
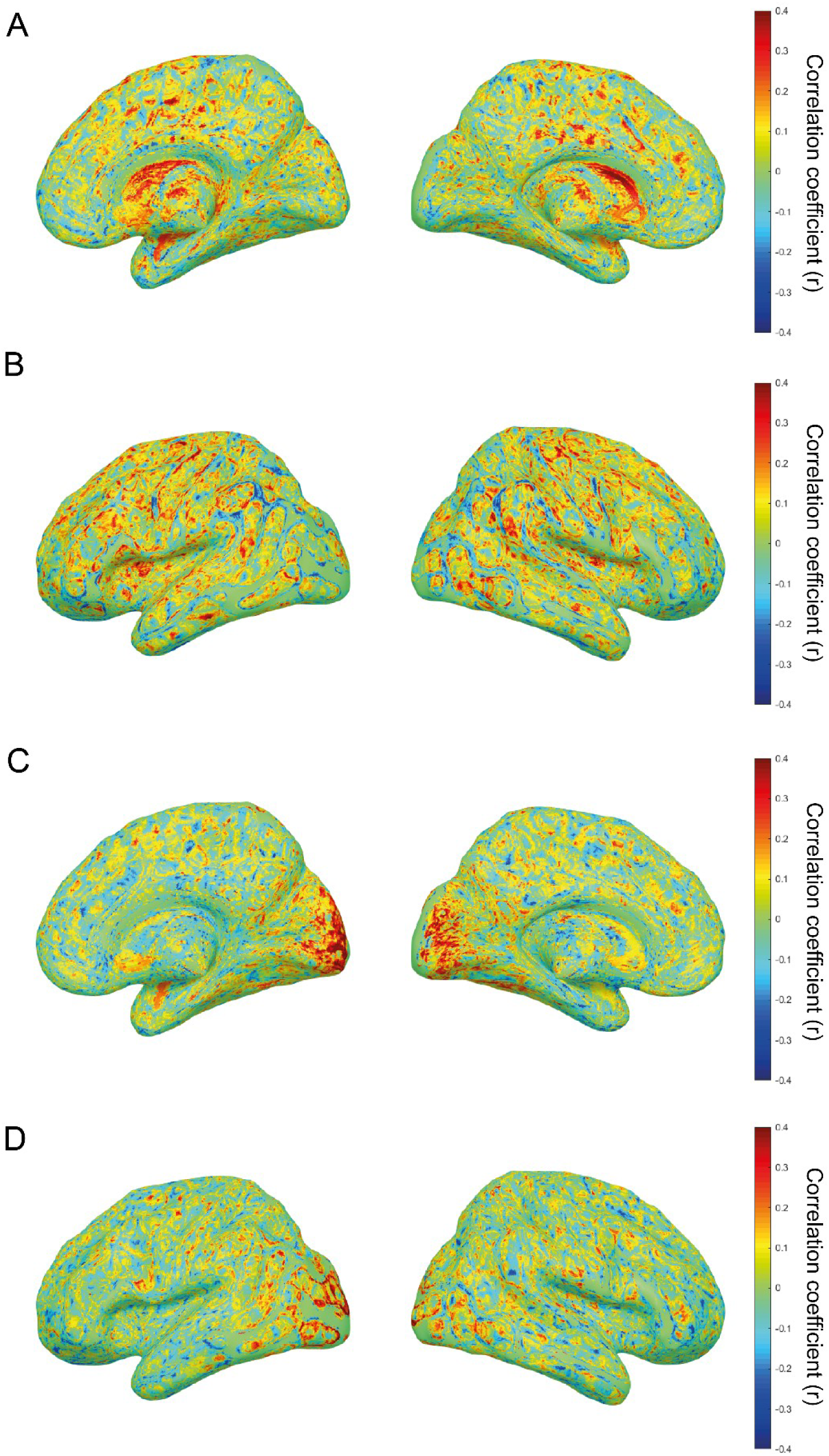
Correlation coefficient (Pearson’s *r*) between the depth of two key regions of interest and the depth of the rest of the brain. The correlation coefficient is shown for left primary motor cortex (a sphere of 8 mm radius centred at MNI coordinates [-37, -25, 62]) in medial (A) and lateral (B) views. The correlation is also shown for left primary visual cortex (sphere centred at [-12, -100, 5]), shown in C and D.

There was no significant correlation between the depth of M1 and the depth of V1 in this sample [r(92)=0.1575, p=0.1295]. The depth of M1 correlated with age in this sample [r(92)=0.3459, p<0.001], but the depth of V1 did not [r(92)=0.0814, p=0.4354]. The depth of M1 was not different between male and female images [t(92)=0.9239, p=0.3579], however V1 was deeper in male images than in female [t(92)=2.7086, p=0.0081, d=0.5670].

## Discussion

The goal of this study was to understand to what extent we can rely on model-based targeting and dose-setting in non-invasive brain stimulation. Or, to put it another way, do we need to have an individual image of each participant’s head in order to be able to set dosage? Since the depth of the target brain area from the scalp surface is a key determinant of the amount of stimulator output required to have an effect on the tissues of the brain, we can frame this as a question about the variance in depth of brain areas in a standardised brain model. A high level of variance in a given area would suggest that that area could not be reliably stimulated without a person-specific image of that region.

Brain volume decreases with increasing age, in this sample and in others (Barnes et al., 2010; Lemaître et al., 2005; Marcus et al., 2007). However the volume enclosed by the skull (here, estimated total intracranial volume) does not decrease. While some of the decrease in brain volume is accounted for by a relative increase in the size of the ventricles, this disparity means that the brain shrinks away from the skull, so increasing distance from the scalp. While the sample included here only covers the middle decades (20s to 60s), it is likely that the distance from scalp to brain surface will change more noticeably in an older sample (Lemaître et al., 2005). Indeed the depth of M1 did correlate with age in this sample, highlighting the need for person-specific models in older populations (Davis, 2017b).

The variance in scalp-to-cortex distance varies across the cortical surface. We might expect the variance to scale with the mean. However the scale-free measure of coefficient of variation is not constant across the cortex. Figures 3 and 4 illustrate this pattern, indicating that some regions of the cortex have a particularly high level of between-subject variance. Analysing a set of functionally important regions of interest suggest that these different areas differ in variance, with relatively high coefficient of variance in the occipital lobe, and lower variance in prefrontal areas. This finding is consistent with the low reliability of phosphene induction measures that target the occipital cortex (e.g. Stewart et al., 2001).

Given the differing patterns of depth variance across the cortex, are there any areas where the depth can be reliably predicted from the depth of either the primary motor or the primary visual cortex? The correlation of cortical depth across the brain surface is generally low, when taking V1 and M1 as a seed region, as shown in Figure 6. The implication of this low correlation is that to have an effect on a behaviour through stimulation of a non-primary target area, one must increase the output of the stimulator to a high enough level that it will overcome the possible additional depth implied in the wider variance, which means a higher level of stimulation for regions that are shallower. Or, conversely, the selected level of stimulation may under-stimulate the brain, so missing potentially important interventions or treatments.

Previous authors have suggested that for TMS studies, the scalp-to-cortex distance offers a simple metric for scaling the intensity of stimulation. Stokes and colleagues (Stokes et al., 2013; Stokes et al., 2005; Stokes et al., 2007) found a pleasingly linear relationship between the scalp-to-cortex distance and TMS-induced motor threshold, with the stimulator intensity required to induce a visible twitch increasing by 2.9% for each millimetre of distance. The results of the present study broadly agree with those of Stokes and colleagues, with similar relative mean depths across brain areas. However these works did not explore the variance of these scalp-to-cortex measures. The relative variance of depth of different brain areas affects the utility of Stokes and colleagues’ simple scaling parameter, as a brain area that shows high inter-subject variability will need more information than simply the mean depth.

Some possible limitations of this study arise from the brain image data. One potential problem is that the sample of participants were recruited from the local area of Washington University (Marcus et al., 2007). The racial demographics of the participants are not reported by Marcus et al., but it would be reasonable to assume that they reflect the demographics of the local area. However the race of the person matters for non-invasive brain stimulation (e.g. for tDCS: Datta, Thomas, Huang, & Venkatasubramanian, 2018), so the findings here may not generalise well. Additionally, the data here was deliberately selected to exclude people with a diagnosis of dementia, but it is possible that the dataset includes people with other disorders that affect neural structure, but which were not identified by the otherwise comprehensive screening performed by Marcus and colleagues. As well as these factors, only a single method of analysis and a single analysis package were used here (namely, SPM 12b). The findings may therefore be sensitive to the analysis pipeline, as is found in other neuroimaging contexts (Poldrack et al., 2017). Finally, the measure of CoV is primarily a descriptive measure. While some fields have attempted to define acceptable limits for CoV, these have tended to be unachievably low for NIBS purposes (e.g. single-digit CoV in veterinary science: Harr, Flatland, Nabity, & Freeman, 2013), where one study has shown CoV of around 70% even for neuronavigated MEPs (Julkunen et al., 2009); the field of brain stimulation needs an understanding of sources and propagators of variance, and their acceptable limits.

Non-invasive brain stimulation has a clear role to play in treating neurally-mediated disorders. TMS and tDCS are very cheap at the point of delivery, meaning that an investment in stimulation equipment may create an opportunity for accessible treatment in a variety of settings. This relative cheapness means that treatment with NIBS may be an attractive option in developing areas where drug treatments may be beyond reach. While the bioethics of NIBS has been explored by several authors in recent years (e.g. Davis, 2017a; Maslen, Douglas, Cohen Kadosh, Levy, & Savulescu, 2014; Wexler, 2015), these discussions have tended to focus on the principles of beneficience (benefit), maleficience (harm) and issues of autonomy; the cheapness of the equipment tending to make access issues (justice) less relevant. However, this study has shown that equipment costs are only a small part of the overall cost of treatment with brain stimulation, as to deliver an effective treatment it is necessary also to have a structural image of the brain. This adds a large cost to an otherwise cheap treatment, and raises the possibility that effective treatment with NIBS may be beyond reach for some healthcare providers.

The present study has shown that for most areas of the cortical surface the variance in depth of the target area is large. The implication of this finding is that studies that use non-invasive brain stimulation, but which do not use person-specific neuronavigation, risk adding a degree of uncertainty to the effects of stimulation, through uncertainty about the depth at which their target lies.

## Author Contributions

NJD conceived the study, wrote the code, and wrote the manuscript.

## Data Accessibility Statement

The original data in this work is available from the Open Access Series of Imaging Studies (OASIS): https://www.oasis-brains.org/. All original computer code is freely available at https://github.com/nickjdavis/ScalpGM

## Abbreviations

TMS: transcranial magnetic stimulation;
tDCS: transcranial direct current stimulation.

